# Repressing PTBP1 is incapable to convert reactive astrocytes to dopaminergic neurons in a mouse model of Parkinson’s disease

**DOI:** 10.1101/2021.11.12.468309

**Authors:** Weizhao Chen, Qiongping Zheng, Qiaoying Huang, Shanshan Ma, Mingtao Li

**Affiliations:** Guangdong Provincial Key Laboratory of Brain Function and Disease, Zhongshan School of Medicine, Sun Yat-sen University, Guangzhou, Guangdong, China, 510080; Department of Pharmacology, Zhongshan School of Medicine, Sun Yat-sen University, Guangzhou, Guangdong, China, 510080

**Author notes:** Indicates equal contribution (Weizhao Chen, Qiongping Zheng). To whom correspondence should be addressed at (S. Ma), (M. Li).

**Keywords:** lineage reprogramming, astrocyte-to-iDAn conversion, PD, PTBP1, AAV leakage, knockdown, conditional deletion, lineage tracing, quiescent astrocyte, reactive astrocyte, 6-OHDA model

## Abstract

Lineage reprograming of resident glia cells to induced dopaminergic neurons (iDAns) holds attractive prospect for cell-replacement therapy of Parkinson’s disease (PD). Recently, whether repressing polypyrimidine tract binding protein 1 (PTBP1) could truly achieve efficient astrocyte-to-iDAn conversion in substantia nigra and striatum aroused widespread controversy. Although reporter positive iDAns were observed by two groups after delivering adeno-associated virus (AAV) expressing a reporter with shRNA or Crispr-CasRx to repress astroglial PTBP1, the possibility of AAV leaking into endogenous DAns could not be excluded without using a reliable lineage tracing method. By adopting stringent lineage tracing strategy, two other studies showed that neither knockdown nor genetic deletion of quiescent astroglial PTBP1 fails to obtain iDAns under physiological condition. However, the role of reactive astrocyte might be underestimated since upon brain injury, reactive astrocyte could acquire certain stem cell hallmarks which may facilitate the lineage conversion process. Therefore, whether reactive astrocytes could be genuinely converted to DAns after PTBP1 repression in a PD model needs further validation. In this study, we used *Aldh1l1-CreER*^*T2*^-mediated specific astrocyte-lineage tracing method to investigate whether reactive astrocytes could be converted to DAns in the 6-OHDA PD model. However, we found that no astrocyte-originated DAn was generated after effective knockdown of astroglial PTBP1 either in the substantia nigra or in the striatum, while AAV “leakage” to nearby neurons was observed. Our results further confirmed that repressing PTBP1 is unable to convert astrocytes to DAns no matter in physiological or PD-related pathological conditions.

**Graphic abstract:** 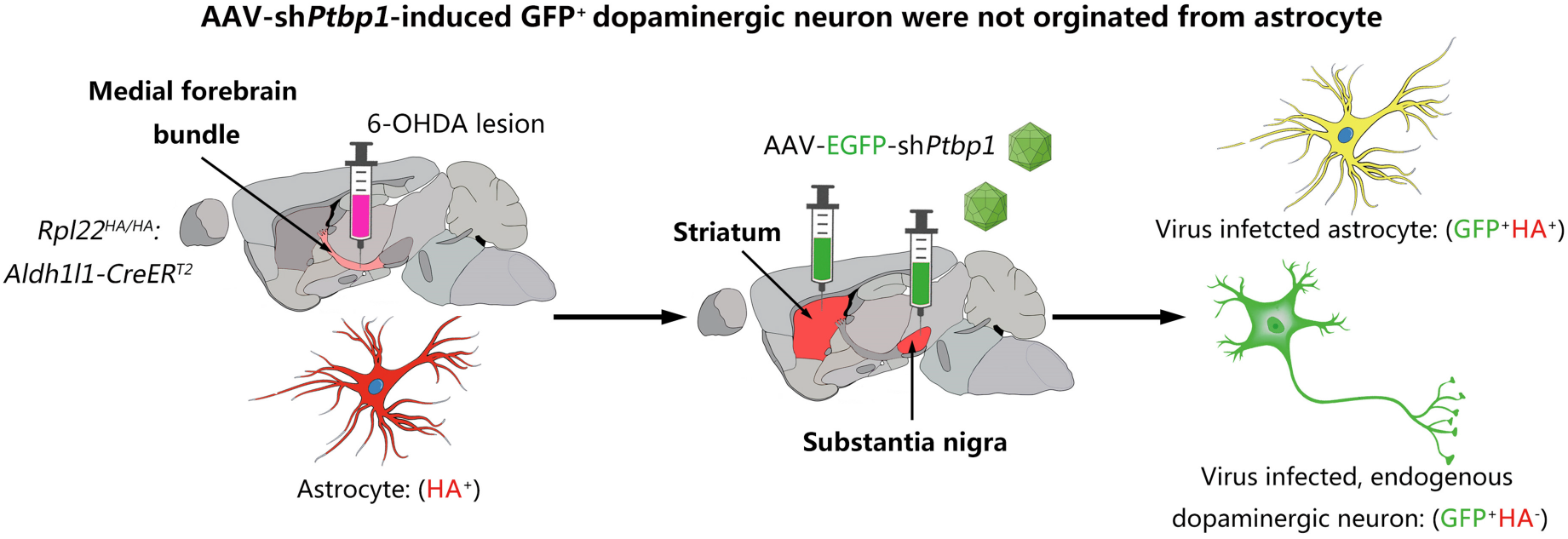

**Highlights:**

- AAV-sh*Ptbp1* rapidly and efficiently induces viral-reporter-labeled DAns in mouse brain under physiological condition
- Viral-reporter-positive DAns are not originated from PTBP1 repressed and lineage traced reactive astrocytes in a mouse PD model

## INTRODUCTION

The emergence and fast development of *in vivo* cell reprogramming technology converting deleterious astrocytes to functional neurons holds great promise for neuroregenerative therapy(Torper and Gotz, 2017). Various groups around the world had successfully achieved astrocyte-to-neuron (AtoN) conversion by forced expression of different pro-neural transcription factors (TF) such as Ngn2(Grande et al., 2013), Ascl1(Liu et al., 2015), NeuroD1(Guo et al., 2014), SOX2(Niu et al., 2013) or various TF combinations(Rivetti di Val Cervo et al., 2017; Wu et al., 2020; Lentini et al., 2021). Differing from the TF over-expression approach, through repressing an RNA-binding protein-PTBP1, two groups respectively reported that functional neurons including DAns could be induced from astrocyte rapidly and efficiently *in vivo*, reconstructing the nigrostriatal circuit and improving motor deficit in a mouse PD model(Qian et al., 2020; Zhou et al., 2020).

Nevertheless, without substantiating the exact origin of the nascent iDAns using reliable lineage tracing strategy, these two outstanding works soon aroused widespread debate and argument (Arenas, 2020; Jiang et al., 2021; Qian et al., 2021). Most recently, by adopting stringent lineage tracing method, two studies arguing against previous findings were published. One group showed that AAV-sh*Ptbp1*-induced, presumed astrocyte-converted iDAns were not truly transdifferentiated from astrocytes but merely AAV-infected endogenous neurons due to virus leakage(Wang et al., 2021). Another group reported that no astrocyte-derived neuron including DAn was generated in multiple brain regions including substantia nigra and striatum in astrocyte-specific *Ptbp1* deletion mice. (Blackshaw et al., 2021). However, both studies simply focused on quiescent astrocytes instead of reactive astrocytes and whether reactive astrocytes could be converted to neurons more effectively after PTBP1 repression needs further verification.

During brain injury or neurodegenerative disease, astrocytes become activated and acquire certain characteristics of neural stem cells (NSCs) such as proliferative, Nestin- or Vimentin-immunoreactive and even multipotent(Buffo et al., 2008; Robel et al., 2011; Shimada et al., 2012; Sirko et al., 2013). Some researchers further claimed that reactive astrocytes with stem cell hallmarks could be reprogrammed to neurons more easily and more efficiently than quiescent astrocytes(Grande et al., 2013; Guo et al., 2014; Wan et al., 2014; Brulet et al., 2017; Mattugini et al., 2019). Therefore, we adopted 6-hydroxydopamine (6-OHDA) PD model with lineage tracing method to investigate whether reactive astrocytes could truly be converted to neurons including DAns.

## Materials and Methods

### Animals

All animal experiments were performed in accordance with the guidelines of the Institutional Animal Care and Use Committee of University. The protocol was reviewed and approved by the Ethics Committee on Laboratory Animal Care. The mice were housed in rooms with controlled 12 hours light/dark cycles, temperature, and humidity, and food and water were provided ad libitum. Eight- to ten-week-old C57BL/6 mice weighing 22–26g were obtained from the Beijing Vital River Laboratory Animal Technological Company (Beijing, China). *Aldh1l1-CreER*^*T2*^ transgenic mice and *Rpl22*^*HA/HA*^ mice (Ribotag) were obtained from The Jackson Laboratory (Stock number #029655, #011029). Hemizygous *Aldh1l1-CreER*^*T2*^ males or females were used for breeding to the *Rpl22*^*HA/HA*^ mice. *Aldh1l1-CreER*^*T2*^: Ribotag mice aged at 8-10 weeks were used for lineage tracing experiments.

### Tamoxifen(TAM) administration

The protocol of TAM administration was determined according to previous work(Srinivasan et al., 2016) with little modifications. Briefly, TAM free-base (Sigma, China) was dissolved in corn oil (Aladdin, China) at a concentration of 10 mg/mL in a 60°C water-bath for 30 minutes. TAM was orally administered at a daily dose of 100 mg/kg body weight for five consecutive days. Experiments were performed two weeks after the last TAM administration.

### 6-OHDA model

The procedure was based on previous study with few modifications(Rivetti di Val Cervo et al., 2017; Qian et al., 2020; Zhou et al., 2020). In brief, 6-OHDA (Sigma, USA) was dissolved in ice-cold saline solution (0.9% NaCl) containing 0.2 mg/mL L-Ascorbic acid (BBI Life Sciences, China) at a concentration of 3 mg/mL. Mice were anesthetized with 3% isoflurane and then placed in a stereotaxic instrument (Model 940, Kopf Instruments, USA). After anesthesia, mice were delivered with 1 μL of 6-OHDA solution (3 mg) into the right medial forebrain bundle (mFB) at 100 nL/min according to the following coordinates: anteroposterior (A/P) = -1.20 mm, mediolateral (M/L) = 1.30 mm, dorsoventral (D/V) = -4.75 mm. Injections were conducted with a 10 μL syringe connected to a 33-Ga needle (Hamilton, USA) using a microsyringe pump (KDS LegatoTM 130, USA). After 6-OHDA injection, mice were typically allowed to recover for three weeks with intense daily care.

### AAV production and infection

AAV2/5-*hGFAP*-EGFP-5’miR-30a-shRNA(*Ptbp1*)-3’miR-30a-WPREs (AAV-sh*Ptbp1*, 3.41 × 10^12^ vg/mL) and the control virus AAV2/5-*hGFAP*-EGFP-5’miR-30a-shRNA(scramble)-3’miR-30a-WPREs (AAV-shscramble, 2.57 × 10^12^ vg/mL) were synthesized based on the pAAV-*hGFAP*-EGFP-WPRE-hGH plasmid (Addgene # 105549) and packaged by Brain VTA (Wuhan, China). For PTBP1 repression, we used the same mouse *Ptbp1* target sequence(5’-GGGTGAAGATCCTGTTCAATA-3’) as previously reported (Qian et al., 2020) while AAV-shscramble expressing scramble shRNA (same nucleotide composition but not in a same sequence order) was used as control.

Before injection into the mouse brain, the AAVs were adjusted to 1 × 10^12^ vg/mL using sterile Dulbecco’s Phosphate Buffered Saline (DPBS, Gibco, USA). Three weeks after 6-OHDA lesion, mice were subjected to AAV injection into the substantia nigra (1 μL) or the striatum (2 μL), respectively, at a speed of 100 nL/min. The coordinates indicating distance (mm) from bregma were A/P= -2.90 mm, M/L= 1.30 mm, and D/V= -4.35 mm for the substantia nigra, and A/P= 0.80 mm, M/L= 1.60 mm, and D/V= -2.80 mm for the striatum. After injection, the needle remained in place for at least five minutes to prevent retrograde flow along the needle track and the needle was slowly removed from the mouse brain. Cleaning and suturing of the wound were performed after the needle was removed.

### Immunofluorescent analysis

For immunofluorescent analysis, mice were anesthetized with 1.25% Avertin and were perfused intracardially with ice-cold phosphate buffered saline (PBS), followed by 4% paraformaldehyde (PFA, Sigma, China) in PBS at a flow rate of 10 mL/min. The brains were then removed and post-fixed in 4% PFA at 4°C overnight (8-12 hours), followed by immersion in 20% and 30% sucrose for 24 hours respectively. Immunofluorescent analysis was performed as previously described (Yu et al., 2018; Hu et al., 2019). In brief, cryostat-coronal sections encompassing the entire midbrain (20 μm) and the striatum(30 μm) were serially collected. Free-floating sections were pre-incubated in blocking solution containing 5% normal donkey serum and 0.3% Triton X-100 in 50 mM Tris-buffered saline (TBS, pH = 7.4) at room temperature for 1 h. Primary antibodies against HA tag (Rabbit, Abcam, ab9110, 1:1000), TH (Chicken, Millipore, AB152, 1:1000), NeuN (Mouse IgG1, Millipore, MAB377, 1:1000) and PTBP1 (Rabbit, Invitrogen, PA581297, 1:1000) were dissolved in diluent and incubated with sections overnight at 4°C. After washing three times, sections were incubated with the secondary antibodies (Thermo Fisher or Jackson ImmunoResearch), which were conjugated with Alexa 488, Alexa 555 or Alexa 647 at room temperature for 1 h. Finally, the sections were visualized under a confocal laser scanning microscope (LSM 780, Carl Zeiss, Germany).

### Statistics

GraphPad Prism version 8.0 was used for statistical analysis. All data were presented as mean ± SEM. Paired-samples t test was used to determine the statistical significance (*p* < 0.05).

## Results

### Repressing PTBP1 rapidly and efficiently induces viral-reporter-labeled neurons

To effectively repress astroglial PTBP1 *in vivo*, we designed and synthesized AAV (serotype 2/5) expressing EGFP followed shRNA targeting mouse *Ptbp1* as reported(Qian et al., 2020), under *hGFAP* promoter (AAV-sh*Ptbp1*), which aimed for specific astrocytes targeting. A corresponding virus expressing scramble shRNA (AAV-shscramble) was used as control (Fig.1A).

**Fig. 1.**
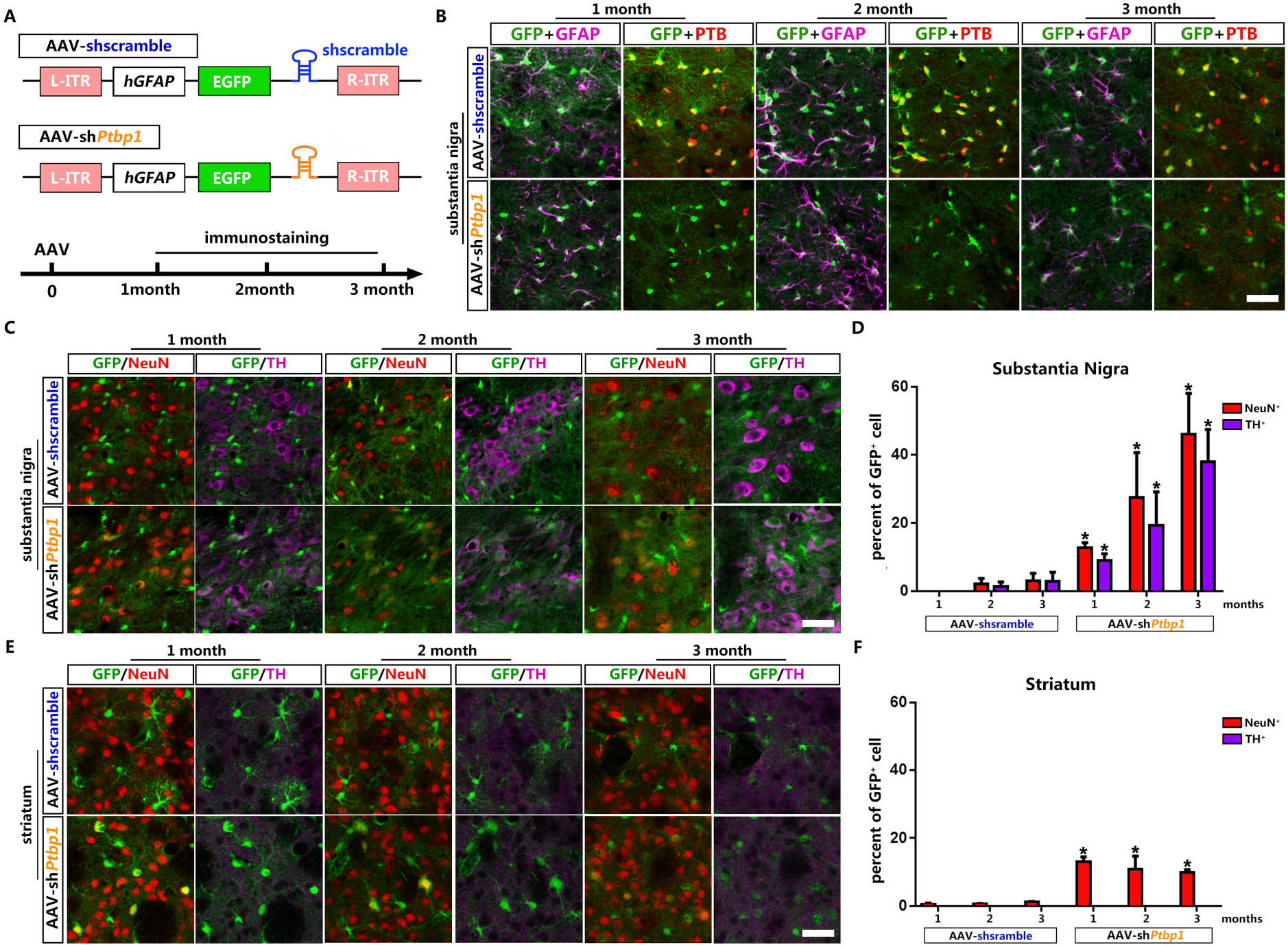
Reporter^+^ neurons were gradually induced in the substantia nigra and the striatum after astroglial PTBP1 repression by AAV-sh*Ptbp1*. (**A**) Schematic of AAV-sh*Ptbp1* and AAV-shscramble vector design and the experimental design. (**B**,**C**,**E**) Representative images of brain slices co-stained GFP (green) with GFAP (purple) or PTBP1 (red) in (**B**), with NeuN (red) or TH (purple) in (**C** & **E**) at indicated time points after AAV-sh*Ptbp1* or AAV-shscramble delivery. Scale bar, 50μm. (**D** & **F**) Number of GFP^+^ cells in the substantia nigra (**D**) and in the striatum (**F**) that show positive staining for NeuN (red) and TH (purple). n = 3 biological repeats. Data are presented as mean ± SEM. *Indicates significant difference between AAV-sh*Ptbp1* and AAV-scramble.

In order to investigate whether repressing astroglial PTBP1 could gradually convert astrocytes to DAns in the substantia nigra and the striatum, brain slices of different time points (1, 2 and 3 months) after AAV injection were collected for analysis. PTBP1 expression was not affected by AAV-shscramble (Fig. 1B upper panel) while it was downregulated to undetectable level by AAV-sh*Ptbp1* in GFAP^+^GFP^+^ cells (Fig. 1B lower panel) from 1 month to 3 months, indicating astroglial PTBP1 was consistently repressed.

Pan-neuronal marker Neuronal Nuclei (NeuN) and DAn marker Tyrosine Hydroxylase (TH) were then co-stained with GFP respectively. The results showed that very few GFP^+^NeuN^+^ cells (approximately 1-2%) were detected even at 3 months after AAV-shscramble injection, while remarkable GFP^+^NeuN^+^ cells were detected, with 12%, 27% and 46% GFP^+^ cells expressed NeuN, among them 9%, 20% and 28% expressed TH (Fig. 1C), at 1, 2 and 3 months after AAV-sh*Ptbp1* injection in the substantial nigra. The gradual increase in numbers of GFP^+^NeuN^+^ cells in AAV-sh*Ptbp1* injected in striatum was similar to that in the substantial nigra. But no GFP^+^TH^+^ cell was found in the striatum. These results highly resemble Fu’s study(Qian et al., 2020) but against Yang’s study(Zhou et al., 2020).

However, all these results are not sufficient to prove that astrocytes were truly converted to neurons or DAns as recently reported AAV-mediated gene expression leakage into neuron(Wang et al., 2021).Thus, more solid evidence is needed to verify the exact origin of the viral-reporter-labeled neurons.

### PTBP1 repression fails to convert quiescent astrocytes to DAns

Genetic lineage tracing(Kretzschmar and Watt, 2012) has been widely recognized as the most convincing strategy for cell source identification, generally performed by combining a cell-specific Cre recombinase expressing mice with a Cre-activated reporter mice. *Aldh1l1-CreER*^*T2*^ mice with highest specificity to target astrocytes(Srinivasan et al., 2016) were chosen to cross bred with a reporter mice *Rpl22*^*HA/HA*^ (Ribotag)(Sanz et al., 2009), whose endogenous ribosomal protein Rpl22 is tagged with three copies of the hemagglutinin (HA) epitope after Cre-mediated recombination (Fig. 2A). After TAM-mediated induction of CreER^T2^ activity, over 99% Aldolase C (AldoC) positive astrocytes were specifically labeled with HA epitope (Fig. 2B) and almost no HA leaky expression in neuron in the substantial nigra and the striatum of *Aldh1l1-CreER*^*T2*^:Ribotag mice.

**Fig. 2.**
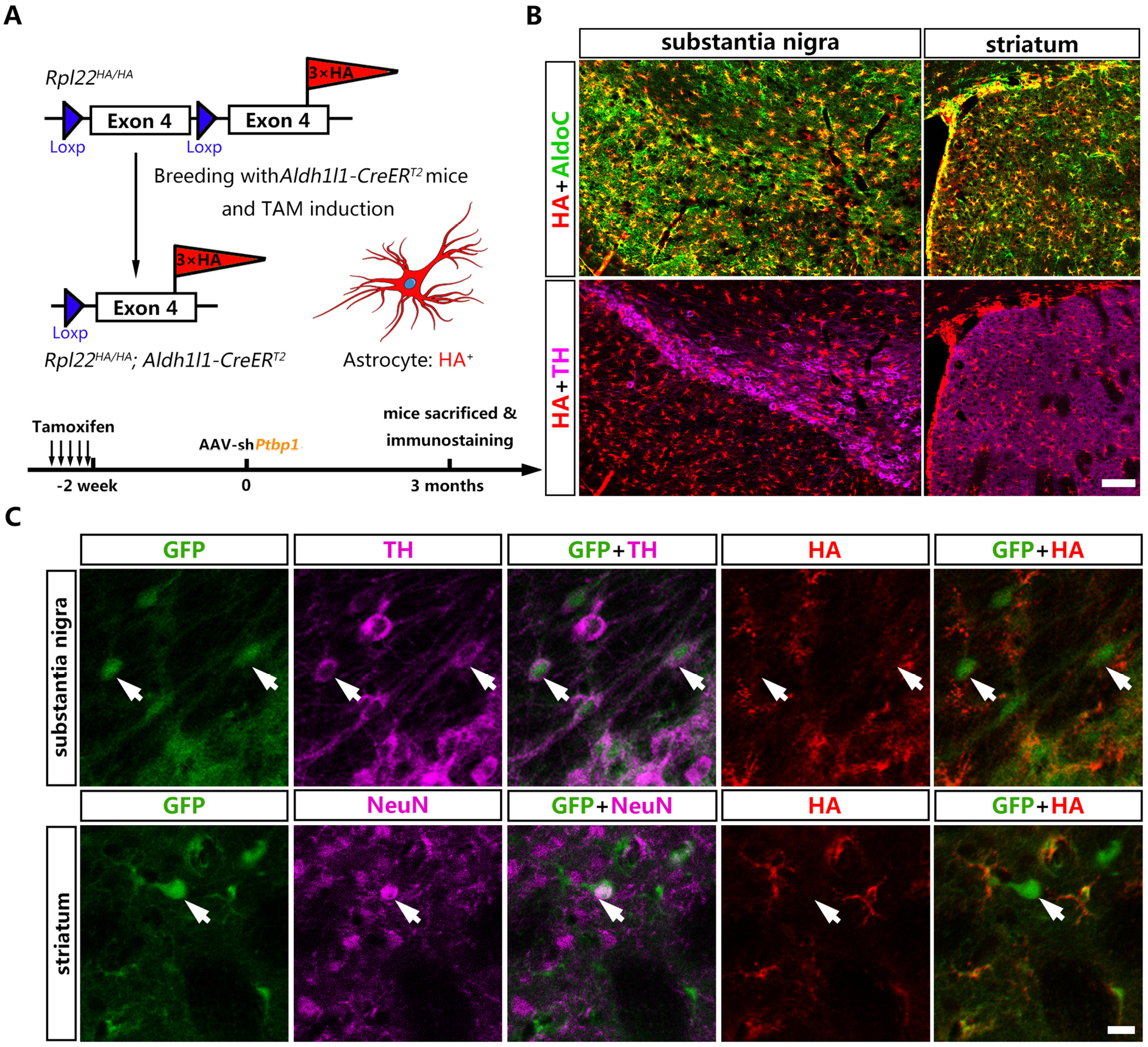
AAV-sh*Ptbp1*-induced reporter^+^ neurons including DAns were not derived from HA^+^ quiescent astrocytes. (**A**) Schematic of breeding strategy of *Aldh1l1-CreER*^*T*2^:Ribotag mice and experimental design. (**B**) Representative images of substantia nigra or striatum of *Aldh1l1-CreER*^*T*2^:Ribotag mice co-stained HA (red) with pan-astrocyte marker AldoC (green) and TH (purple) 2 weeks after TAM administration. Scale bar, 100μm. (**C**) Representative images of brain slices co-stained GFP (green), HA (red) with TH (purple) in the substantia nigra and with NeuN (purple) in the striatum 3 months after AAV-sh*Ptbp1* delivery. Scale bar, 75μm.

Two weeks after TAM induction, *Aldh1l1-CreER*^*T2*^:Ribotag mice were injected with AAV-sh*Ptbp1* into the substantia nigra or the striatum to verify whether the GFP^+^TH^+^ or GFP^+^NeuN^+^ cells were originated from HA-labeled astrocytes. Three months later, mice were sacrificed for triple immunostaining of GFP, HA and NeuN or GFP, HA and TH. Through exhaustive examination of the whole midbrain and striatum, we couldn’t find any GFP^+^TH^+^ or GFP^+^NeuN^+^ cells that were simultaneously HA-positive(Fig. 2C), suggesting they were not converted from astrocytes but probably endogenous neurons infected with AAV and expressing GFP reporter.

Therefore, the above results clearly illustrated that PTBP1 repression fails to convert quiescent astrocytes to neurons including DAns, which is consistent with recent studies(Blackshaw et al., 2021; Wang et al., 2021).

### PTBP1 repression also fails to convert reactive astrocytes to DAns in a 6-OHDA model

A lot of evidence has suggested that reactive astrocytes may acquire certain characteristics of NSC upon brain injury, which could promote AtoN conversion(Grande et al., 2013; Guo et al., 2014; Wan et al., 2014; Brulet et al., 2017; Mattugini et al., 2019). To verify whether repression PTBP1 could converts reactive astrocyte to neuron including DAns, we performed 6-OHDA lesion in the mFB on *Aldh1l1-CreER*^*T2*^:Ribotag mice 2 weeks after TAM administration.

Three weeks after 6-OHDA lesion in mFB, the mice were subjected to AAV-shscramble or AAV-sh*Ptbp1* injection in the substantia nigra and the striatum respectively, and then sacrificed for immunostaining after three months (Fig. 3A).

**Fig. 3.**
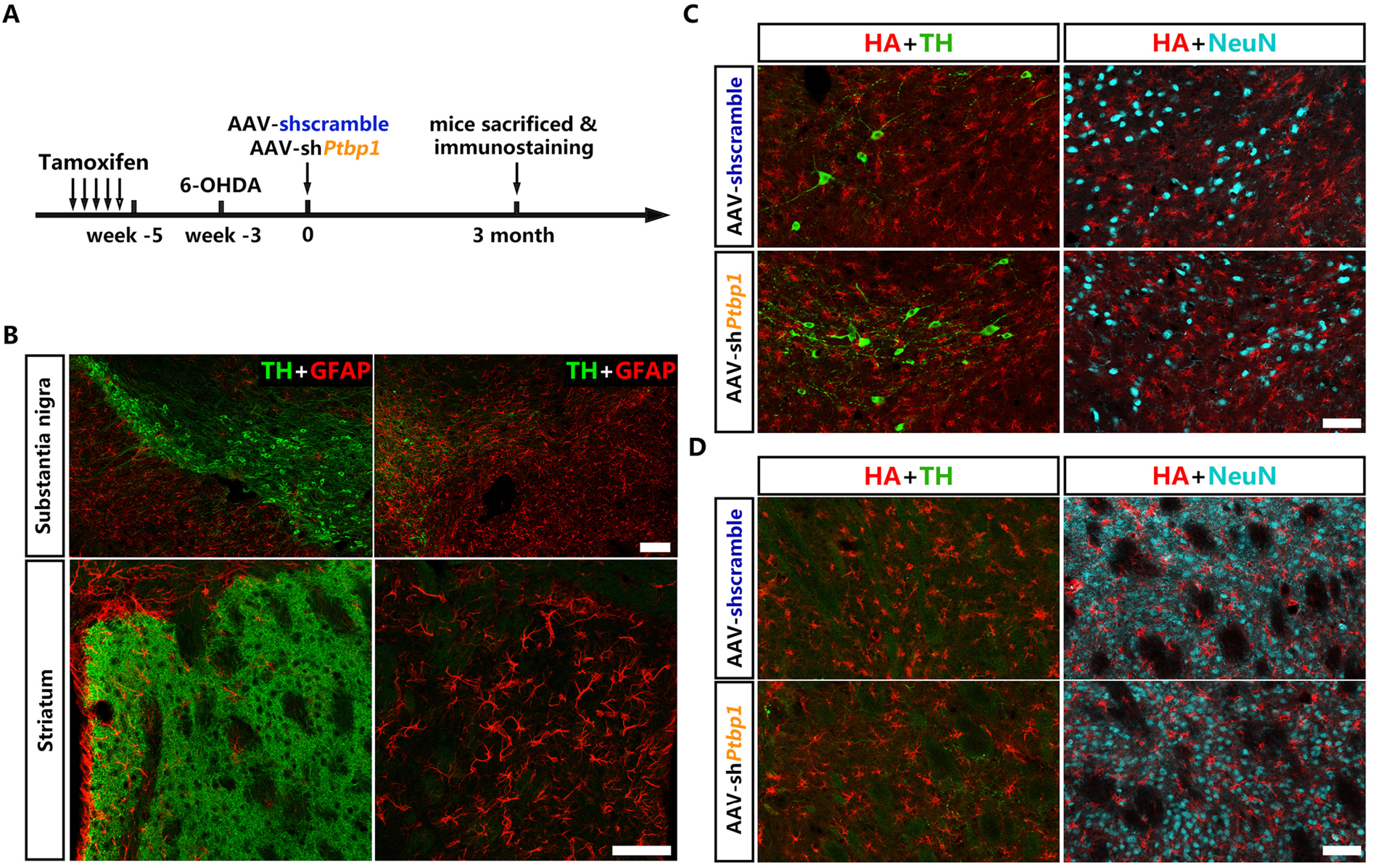
No neuron including DAn co-staining HA can be detected after astroglial PTBP1 repression in 6-OHDA model. (**A**) Schematic of experimental design. (**B**) Representative images of substantia nigra or striatum after 6-OHDA lesion co-stained with TH (green) and GFAP (red). Scale bar, 100μm for the substantia nigra and 50μm for the striatum. (**C**,**D**) Representative images of HA (red), TH (green) and NeuN (cyan) co-stained substantia nigra (**C**) and striatum (**D**) 3 months after AAV-sh*Ptbp1* or AAV-shscramble delivery in 6-OHDA model. Scale bar, 25μm.

Our results showed that 6-OHDA induced severe lesion to the nigrostriatal pathway with significantly reduced the number of DAns in substantia nigra and the densities of TH^+^ fibers in the striatum. Meanwhile, astrocytes become remarkably activated, indicated by classic cytoskeletal and morphological changes including hypertrophy of the main processes and cell body and upregulation of intermediate filament proteins such as GFAP (Fig. 3B). However, no HA^+^ cells co-expressing NeuN or TH could be detected either in the substantia nigra or the striatum (Fig. 3C & D),no morphologic change of astrocytes could be seen after PTBP1 repression (Fig. 3C & E) and only subtle changes in gene expression could be seen after astroglial *Ptbp1* deletion(Blackshaw et al., 2021), suggesting that neither AtoN nor astrocyte-to-DAn conversion occurred. Together, these data demonstrate that repressing PTBP1 is also incapable to generate DAns from reactive astrocytes in a murine PD model.

In summary, we provide solid and convincing evidence that repressing astroglial PTB using shRNA-based RNA interference (RNAi) technology is unable to generate DAns from astrocytes neither in quiescent nor in reactive state.

## Discussion

In summary, we provide solid and convincing evidence that repressing astroglial PTBP1 using shRNA-based RNA interference (RNAi) technology is unable to generate DAns from astrocytes neither in quiescent nor in reactive state.

### Factors that lead to artificial AtoN conversion

It is well known that excessive dosage of AAV increases the tendency of virus leakage to other cells nearby. Once the astrocyte-specific labeling reporter leaks to neuron, the reporter^+^ neuron would then be misidentified as neuron converted from astrocyte, resulting false-positive results. In addition to virus dosage, other factors that may promote the artificial AtoN conversion in a previous study(Qian et al., 2020) will be discussed below.

Firstly, it is widely accepted that *Gfap* promoter, either employed in transgenic mice or in virus, is not stringent for astrocyte specific labeling and genetic manipulation. Various studies reported that a large proportion of neuron also has *Gfap* promoter activity especially after brain injury(Hol et al., 2003; Su et al., 2004; Lee et al., 2006; Lee et al., 2008; Fujita et al., 2014). Some researchers explained that overdosed AAV would exert toxic effect on neurons(Khabou et al., 2018; Xiong et al., 2019; Xiang et al., 2021) which may lead to a time-dependent activation of the *Gfap* promoter. Alternatively, enhanced activity of *Gfap* promoter may indicate that neuron reverse to an immature transcriptome state upon injury in order to facilitate the neural repair and regeneration process(Poplawski et al., 2020). Moreover, why control virus (AAV-shscramble) seem highly specific to astrocytes but AAV-sh*Ptbp1* that show severe virus leakage to neuron? A recent study explained that the promoter specificity of *Gfap* is likely cis-regulated by NeuroD1 or some other pro-neural TFs(Wang et al., 2021) Therefore, the control virus may no longer be suitable to serve as a control and thus reliable lineage tracing is indispensable for *in vivo* cell reprogramming research.

Secondly, the Cre-activated Loxp-STOP-Loxp (LSL) structure would have leakage expression without Cre recombinase. Even classical LSL structure which has 3-4 repeats of SV40-polyA signal between two Loxp sites can’t avoid leakage(Sohal et al., 2009), not to mention the non-classical Loxp-STOP-Loxp (LSL) structure that only has one SV40-polyA signal located in the ORF of Neo/Kana resistance(Qian et al., 2020).

Therefore, without a reliable lineage tracing method, most of the presumed astrocyte-converted neurons were merely endogenous neurons misidentified by unexpected labeling due to virus leakage. The actual AtoN conversion efficiency is probably highly overestimated.

### Possible reasons for AtoN conversion failure *in vivo*

Compared with the convincing AtoN result *in vitro*(Qian et al., 2020), why PTBP1 repression fail to convert astrocyte to neuron *in vivo*? We suppose that AtoN conversion did not even initiate because the no morphologic change of astrocyte could be seen after PTBP1 repression (Fig. 3C & E) and only subtle changes in gene expression could be seen after astroglial *Ptbp1* deletion(Blackshaw et al., 2021). The reason why the PTBP1-repressed astrocyte fail to initiate the conversion process may attribute to astrocyte itself and the local microenvironment.

On one thing, the status of astrocyte may greatly influence AtoN conversion outcome since studies found that reactive astrocytes could be converted to neuron more easily than quiescent astrocyte(Guo et al., 2014; Brulet et al., 2017). However, in 6-OHDA lesion model we could not find any sign of AtoN conversion although astrocytes became intensively activated. Reactive astrocytes are highly heterogeneous and can be divided into A1 subtype (pro-inflammatory) and A2 subtype (anti-inflammatory) according to different disease model(Zamanian et al., 2012; Escartin et al., 2021). Whether age-related decrease in neuronal reprogramming efficiency(Qian et al., 2020) is associated with gradual conversion of astrocytes to a pro-inflammatory phenotype (A1) with aging(Clarke et al., 2018) and whether anti-inflammatory astrocytes (A2) could be converted to neurons more efficiently need further investigation.

For another, the key difference between *in vitro* and *in vivo* is the local microenvironment. Various bioactive molecules either beneficial (neurotrophic factors) or detrimental (inflammatory cytokines) existed in the local microenvironment(Janowska et al., 2019). However, under pathological condition, these secretory factors are usually deleterious (pro-inflammatory cytokines) to the AtoN conversion process and subsequent survival, maturation of the new-born neurons. Unexpectedly, under special pathological condition like ischemia, astrocytes spontaneously initiate a potent neurogenic program after Notch signaling being repressed(Magnusson et al., 2014). Moreover, studies showed that astrocytes from different brain regions have different reprogramming efficiency and different neuron subtype preference (Grande et al., 2013; Liu et al., 2015; Hu et al., 2019; Mattugini et al., 2019). All these findings suggest that the different local environment have different impact on AtoN conversion efficiency and modulating local environment may be critical for enhancing the efficiency of AtoN conversion.

Therefore, only repressing PTBP1 is not enough to initiate cell fate change of astrocyte towards neuron in the mouse brain. The impact of heterogeneity of both astrocyte subtype and local environment on AtoN conversion need further inquiry in the future.

### Possibility of neuron replenishment and behavioral recovery after PTBP1 repression

According Fu’s study, using antisense oligonucleotides (ASOs) targeting *Ptbp1* (*Ptbp1*-ASO), a fraction of tdTomato-labeled cells in *Gfap-creER™*: Rosa-tdTomato mouse were converted to TH^+^ DAns(Qian et al., 2020). If the astrocyte-to-DAn conversion did not happen, then other cell types rather than astrocyte might contribute to the *Ptbp1*-ASO-mediated neuron conversion which may explain the DA neuron number replenishment, striatal TH^+^ fibers density recovery and behavioral improvement in Fu’s study.

The first possible cell source is NSC since PTBP1 is well known to maintain NSC pools(Shibasaki et al., 2013) and is sharply diminished upon neuronal lineage induction(Boutz et al., 2007; Makeyev et al., 2007). Roy Maimon et al reported that using similar *Ptbp1*-ASO, radial glial-like cells and other GFAP-expressing cells could be converted to neurons(Maimon et al., 2021). Another candidate cell source is oligodendrocyte (OL) which was found to have certain *Gfap* promoter activity (Behrangi et al., 2020). Weinberg et al. reported that OLs could also be reprogrammed to functional neurons after PTBP1 repression(Weinberg et al., 2017). Therefore, these potent cell types might contribute to the neuron restoration, functional and even behavioral recovery but need further and detailed investigation in the future.

## Authors Contribution

The manuscript was written with the contributions of all authors. All authors have approved the final version of the manuscript. W. Chen and Q. Zheng conducted most of the experiments. W. Chen and and Q. Huang conducted the literature search and wrote the manuscript. S. Ma and M. Li provided supervision and constructive feedback during manuscript writing. All authors contributed critical revisions to the manuscript.

## Acknowledgements

We thank Qingxing Zhang for his assistance with the animal breeding and genotyping.

